# Learning Genetic Perturbation Effects with Variational Causal Inference

**DOI:** 10.1101/2025.06.05.657988

**Authors:** Emily Liu, Jiaqi Zhang, Caroline Uhler

**Author notes:** Correspondence: {, }. Equal Contributions (Emily Liu, Jiaqi Zhang).

## Abstract

Advances in sequencing technologies have enhanced the understanding of gene regulation in cells. In particular, Perturb-seq has enabled high-resolution profiling of the transcriptomic response to genetic perturbations at the single-cell level. This understanding has implications in functional genomics and potentially for identifying therapeutic targets. Various computational models have been developed to predict perturbational effects. While deep learning models excel at interpolating observed perturbational data, they tend to overfit and may not generalize well to unseen perturbations. In contrast, mechanistic models, such as linear causal models based on gene regulatory networks, hold greater potential for extrapolation, as they encapsulate regulatory information that can predict responses to unseen perturbations. However, their application has been limited to small studies due to overly simplistic assumptions, making them less effective in handling noisy, large-scale single-cell data. We propose a hybrid approach that combines a mechanistic causal model with variational deep learning, termed Single Cell Causal Variational Autoencoder (SCCVAE). The mechanistic model employs a learned regulatory network to represent perturbational changes as shift interventions that propagate through the learned network. SCCVAE integrates this mechanistic causal model into a variational autoencoder, generating rich, comprehensive transcriptomic responses. Our results indicate that SCCVAE exhibits superior performance over current state-of-the-art baselines for extrapolating to predict unseen perturbational responses. Additionally, for the observed perturbations, the latent space learned by SCCVAE allows for the identification of functional perturbation modules and simulation of single-gene knockdown experiments of varying penetrance, presenting a robust tool for interpreting and interpolating perturbational responses at the single-cell level.

**Author summary:** Understanding how genes interact and respond to perturbations is crucial for uncovering the mechanisms of cells and identifying potential ways to treat diseases. Recent advances in sequencing technologies now allow us to measure how individual cells react when specific genes are altered. However, making sense of this complex data requires advanced computational tools. In our work, we address the challenge of predicting how cells respond to potentially new untested genetic perturbations. We noticed that while deep learning models perform well on data measured before, they struggle with making predictions on new cases. On the other hand, models based on biological understanding can, in theory, make better predictions, but they often rely on overly simple assumptions that do not hold with real-world data. We developed a new method that combines the strengths of both approaches. Our model, called SCCVAE, uses knowledge of gene networks together with deep learning to better predict how cells will respond to gene changes. It can simulate new experiments and help identify groups of genes that work together. This tool could be valuable for researchers studying perturbational changes, as well as gene functions and diseases.

## 1 Introduction

Gene-editing technologies provide useful probes for the study of gene regulation in cells [1]. By perturbing individual genes and observing transcriptomic changes, we can disentangle and resolve the downstream effects of these perturbations. These insights facilitate a range of downstream applications, from identifying genes involved in fundamental cellular processes (e.g., [2, 3, 4]) to discovering potential drug targets for therapeutic use (e.g., [5, 6]). There are numerous potential target genes for perturbation. Perturb-seq allows for large-scale exploration by combining high-throughput CRISPR gene editing with single-cell RNA sequencing [7, 8]. Recent advances have further expanded its scale, enabling the collection of data on genome-wide perturbations in millions of cells [9]. Understanding cellular responses to the genetic perturbations introduced through these high-content data is of great importance.

Various computational approaches have been proposed to interpret and predict perturbational effects. One predominant line of work explores the performative powers and inductive biases brought by popular deep learning architectures. For example, Lotfollahi et al. [10] uses a compositional architecture combined with an adversarial network to disentangle perturbational effects from the basal cell states. Yu and Welch [11] utilizes two separate autoencoders to learn perturbation-specific and cell-specific latent representations and employs a normalizing flow to map between these representations. Roohani et al. [12] uses a graph-based network that leverages a gene ontology-based graph to simultaneously learn perturbation embeddings and their corresponding effects. Lopez et al. [13] proposes a modified variational autoencoder with carefully designed noise models, modeling perturbational effects as sparse shifts of these noise distributions. Wu et al. [14] uses a variational autoencoder with a graph attention architecture to encode gene regulations. Several variational-inference-based analyses have also been proposed to interpret the observed perturbations [15, 16]; see Rood et al. [17] for a comprehensive review. Recently, transformer architectures have also been used to learn unsupervised representations of cells and genes, with perturbation prediction being a downstream task [18, 19]. While these methods excel at interpolating observed data, they are prone to overfitting, which may limit their ability to generalize to unseen perturbations. Ahlmann-Eltze et al. [20] observed that simple linear models can outperform sophisticated deep learning models in generalizing to unseen perturbations. However, their approach focuses on pseudo-bulk analysis, whereas this work aims to resolve perturbational effects at the single-cell level.

There are several axes to consider when generalizing to unseen perturbations. *First*, the generalization task may involve extrapolating from single-gene perturbations to combinatorial perturbations that target multiple genes or to single-gene perturbations targeting novel genes, potentially administered at varying penetrance / multiplicity of infection. In this work, we mainly focus on extrapolations to novel single-gene perturbations. *Second*, depending on the task, the principles one uses for generalization may differ. For example, to generalize to combinatorial perturbations, a popular principle is to use additivity to combine the effects (potentially in a latent space) and learn the interactions as residuals (e.g., [12, 21, 22, 23]). To generalize to unseen novel perturbations, two major principles are to use prior knowledge of how the new target relates to observed targets (e.g., [11, 12] reviewed above) or a mechanistic model that specifies how the perturbation propagates according to a gene regulatory network (e.g., [24]). Prominent deep learning models, as discussed above, are mostly prior-knowledge-based, typically relying on gene ontology annotations, which might suffer from poor generalization as such annotations usually capture partial correlations. On the other hand, mechanistic models hold greater potential for extrapolation as they capture regulatory information. Prior mechanistic models usually make assumptions about how perturbations change transcriptomic profiles. For example, Kamimoto et al. [24], Zhang et al. [25] uses a linear causal model based on a learned gene regulatory network; Dibaeinia and Sinha [26] uses a chemical master equation to simulate transcription. However, their applications have been limited to relatively small studies due to simplistic parametric assumptions and/or expensive stochastic differentiation equation simulations.

In our work, we propose a hybrid model that combines a mechanistic causal model with variational deep learning, termed the Single Cell Causal Variational Autoencoder (SCCVAE). The mechanistic causal model captures regulatory information by employing a learned regulatory network and models perturbations as shift interventions that propagate through this network. It is defined on a lower-dimensional space, capturing essential information to reconstruct the entire transcriptomic readout. To address the issue of simplistic parametric assumptions in most mechanistic models, SCCVAE integrates this model into a variational autoencoder. This integration allows SCCVAE to learn and generate rich, comprehensive transcriptomic responses. Our findings demonstrate that SCCVAE excels at generalizing beyond observed perturbations, enabling accurate predictions of unseen single-gene perturbations, and outperforms both standard and state-of-the-art baselines. The mechanistic model specifies the penetrance of the perturbation, allowing simulation of single-gene knockdown perturbations with varying penetrance. As for the observed perturbations, since SCCVAE learns how the perturbation shifts the variables defined by the mechanistic model, we can extract this information to serve as a perturbation representation, which we observe to capture functional perturbation modules.

## 2 Methods

### 2.1 Variational Causal Model for Perturbations

Consider the gene expressions of a cell, denoted by *X*^*p*^, perturbed by a single-gene perturbation (i.e., single-guide RNA) represented by *p*. When the cell is a negative control (e.g., with non-targeting guide RNAs), we denote its gene expressions as *X*^Ø^ or *X* in short with *p* = Ø. Each cell is measured by the expression levels of *m* distinct genes, i.e., *X*^*p*^ ∈ ℝ ^*m* ×1^. We now explain the components of SCCVAE.

#### 2.1.1 Structural causal model

Central to the design of our model is the concept of structural causal models [27, 28]. A *structural causal model* (SCM) is a framework for understanding complex systems by modeling the regulations between causal variables. Formally, a structural causal model is defined as a tuple ⟨*Z, U, F*⟩ consisting of a set of exogenous noise variables *Z*, a set of endogenous random variables *0055*, and a set of functions *F* : *Z* × *U* → *U*. Each SCM is associated with a directed acyclic graph 𝒢, where node *i* in 𝒢 corresponds to an endogenous variable *U*_*i*_ and edges denote causal relationships between variables. For a given *i*, the endogenous variable *U*_*i*_ depends on its associated exogenous noise variable *Z*_*i*_ and its endogenous parents *U*_pa(*i*)_, where *j* ∈ pa(*i*) if there is an edge *j* → *i* ∈ 𝒢. Mathematically, we have

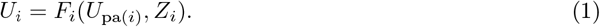

In the context of gene regulations, the endogenous variables *U* ∈ ℝ^*n×*1^ denote an abstracted representation of *n* gene modules, where each *U*_*i*_ corresponds to one gene module. Intuitively, this representation can be interpreted as a low-dimensional feature summarizing the expression of *m* genes in the context of a cell, where *m* can be much larger than *n*. The causal graph *G* indicates gene regulations, where *j* ∈ pa(*i*) if genes in module *j* directly regulate genes in module *i*. The exogenous variables *Z* record the intrinsic variations of *U*, not explained by regulations. Given the values of *U*_pa(*i*)_ and *Z*_*i*_, function *F*_*i*_ then generates *U*_*i*_ associated with module *i*. In our setting, we choose to use a specific parametric family of SCM, where *Z*_*i*_ are assumed to be Gaussian and *F*_*i*_ are linear functions, as this parameterization can encapsulate all multivariate Gaussian distributions, which we observe to sufficiently fit the transcriptomic data. In other words, Eq. (1) is realized by

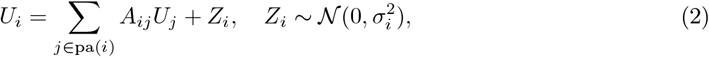

where *A*_*ij*_ describes the contribution of *U*_*j*_ in *U*_*i*_. Since the scaling of the endogenous variables can be set arbitrarily, we further assume that *σ*_*i*_ = 1.

A genetic perturbation *p* alters the endogenous variables *U* into *U* ^*p*^ by modifying the regulatory relationships in Eq. (2). Here, we use an additive shift model, summarized by

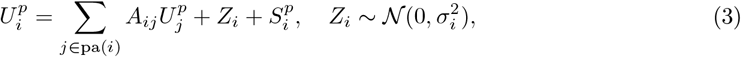

where *S*^*p*^ is the shift vector associated with perturbation *p*. Figure 1 shows an example. In the following section, we explain how to learn this SCM within a variational inference framework.

**Figure 1.**
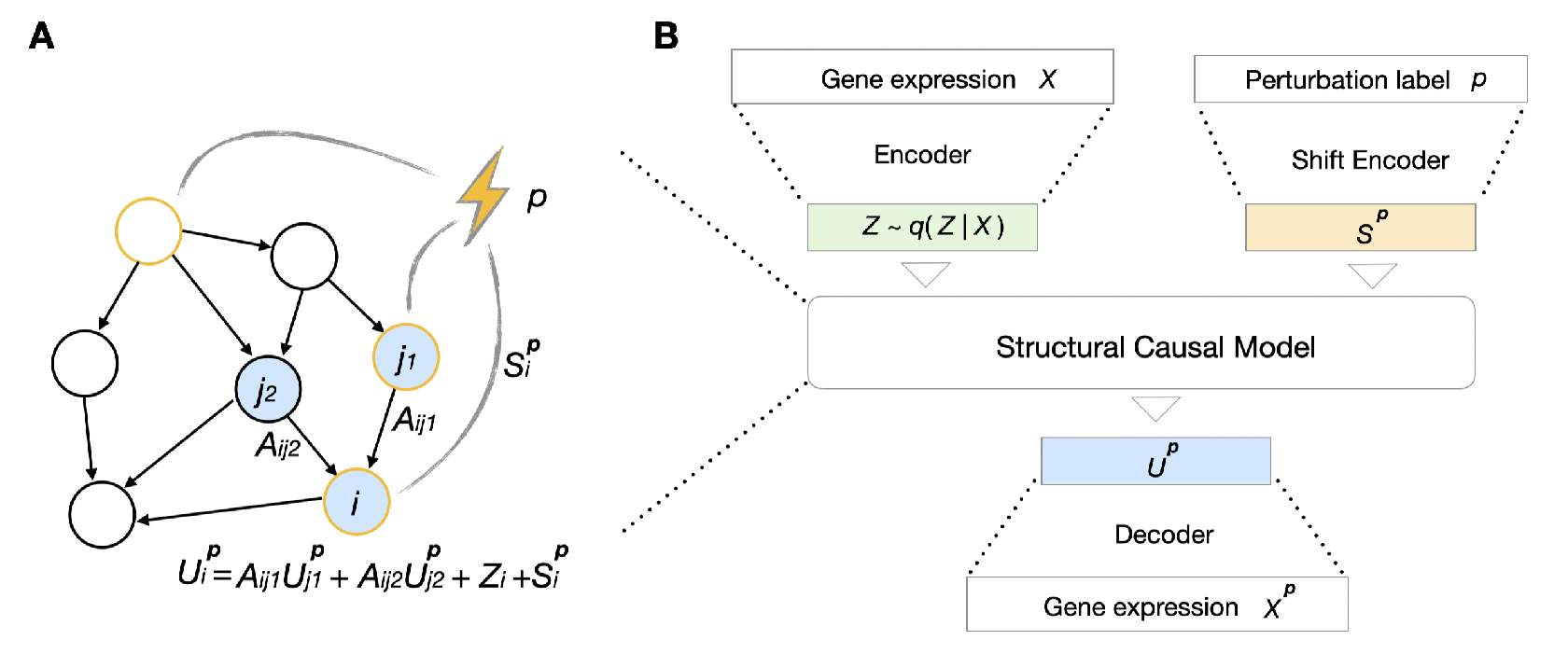
Key components of SCCVAE. (A) Illustration of a structural causal model, where the parameters associated with gene *i* are annotated. (B) The architecture of SCCVAE. It contains: an expression encoder that maps *X* to exogenous noise variables *Z*, a shift encoder that maps *p* to a shift vector *S*^*p*^, a structural causal model (e.g., as illustrated in (A)) that maps *Z, S*^*p*^ to *U* ^*p*^, and an expression decoder.

We make a note here that the definition of SCM assumes acyclic *G* to ensure that Eq. (1) has a unique solution. However, it can be easily extended to cyclic graphs to model feedback loops by setting pa(*i*) as all parent nodes of *i* excluding itself in the cyclic graph.

#### 2.1.2 Hybrid variational causal model

SCCVAE integrates and learns the SCM in a variational autoencoder. It consists of four key components: the *expression encoder*, the *shift encoder*, the *SCM*, and the *expression decoder*. Figure 1 shows all inputs, outputs, and components of the model architecture.

The *expression encoder* takes in the expression of one control cell *X* ∈ ℝ^*m×*1^ and encodes it into the exogenous variables *Z* ∈ ℝ ^*n×*1^, with *n < m*. Intuitively, such *Z* represents intrinsic variations of *n* gene modules, not explained by regulations, that is shared between the control and the perturbed distributions. This set of *n* modules captures essential information to reconstruct the entire transcriptomic readout. In our implementations, we select *n* = 512. We encode the conditional distribution *q*(*Z* | *X*) using diagonal normal distribution, where the mean and variance are parameterized by neural networks, and minimize the KL divergence 𝔼_*X*_ (*D*_*KL*_ (*q*(*Z* | *X*) || *p*(*Z*))), where *p*(*Z*) corresponds to *𝒩* (0, I_*n*_) with I_*n*_ being the identity matrix of order *n*. We then use the reparameterization trick [29] to sample *Z* from *q*(*Z* | *X*) to obtain the endogenous variables.

As inputs to the *shift encoder*, we need a label representing each single-gene perturbation. This label should be attainable for unseen perturbations for generalization purposes. Here we use the top principal components of the transposed control cell expression matrix of shape ℝ^*m×l*^ as perturbation labels, where *l* is the number of control cells. For each of the *m* measured genes, we compute the associated top principal components vector in ℝ^*l*^ using the control cell expression matrix. We take top principal components in this vector as our label representing perturbation on this measured gene. The *shift encoder* encodes a perturbation label *p* into a shift vector *S*^*p*^ ∈ ℝ^*n*×1^. Intuitively, such *S*^*p*^ denotes the direct effects that a perturbation has on these *n* gene modules, capturing both on-target and potentially off-target effects. Such direct effects then propagate to other endogenous variables through Eq. (3). Note that Eq. (2) is a special case of Eq. (3) with *p* = Ø and *S*^*p*^ = 0.

We can stack Eq. (3) for different *i* = 1, …, *n* and write it into a matrix form

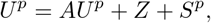

which is equivalent to

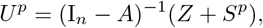

where 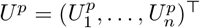 and *A*_*ij*_ = 0 if *j ∉* pa(*i*). Therefore in the *SCM*, we only need to specify the graph *𝒢* and parametrize *A* by masking it according to the sparsity pattern in *𝒢*, upon which *U* ^*p*^ can be directly computed based on *Z* and *S*^*p*^. In our implementations, we find that a learned network *𝒢* (i.e., using an upper-triangular mask) that is optimized during training to work well.

Finally the *expression decoder* takes the perturbed endogenous variables *U* ^*p*^ and constructs the expression profile *X*^*p*^ of perturbed cells. The model is trained using the variational lower bound with an added maximum mean discrepancy (MMD) term [30] specified by

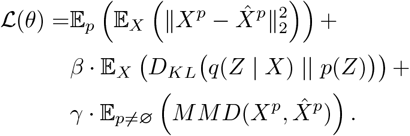

The variational lower bound in the first two terms (with *p* = Ø) maximizes the likelihood of the hybrid causal model over the control distribution, whereas the mean square error in the first term (with *p ≠* Ø) and the added MMD term match and stabilize the training for the perturbational distributions.

Here *θ* subsumes all the unknown parameters, 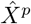 denotes the output of SCCVAE, and hyper-parameters *β, γ* are scaling factors specified in Appendix A.1.

In summation format, 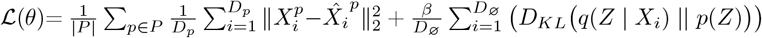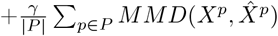, where *P* is the set of all perturbations, *D*_*p*_ is the number of cells for *p* ∈ *P*, and *D*_Ø_ is the number of control cells.

### 2.2 Shift Selection for Unseen Perturbations

When generalizing to an unseen single-gene perturbation, especially in knockdown or activation experiments, it is essential to account for the penetrance of the perturbation, i.e., the magnitude of its effect. This is because such perturbations may be administered with guides of different efficiencies compared to the perturbations seen during training. As a result, simply encoding the identity of the new perturbation is insufficient; we must also infer how strongly it acts. Specifically, for an unseen perturbation *q*, we can encode it into a shift vector *S*^*q*^ ∈ ℝ^*n×*1^ using SCCVAE trained on observed perturbations. To quantify the effects of different penetrance, we attribute one single scalar *c*^*q*^ ∈ ℝ to denote its effect on the endogenous variables. In other words,

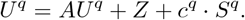

Then *U* ^*q*^ is passed to the decoder to generate in-silico transcriptomic profiles of single cells perturbed by *q*.

If additional metadata exists, e.g., a proliferation screen with the same library, one can use this information to specify the intensity of the perturbation. When such information is not available, one can use, e.g., bulk perturbation data to select *c*^*q*^ in order to evaluate the capability of the model. Specifically, one can grid search for *c*^*q*^ within a range by comparing the average predicted perturbed cell expression 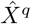 with the bulk expression of cells subjected to perturbation *q* using mean squared error. The accurate penetrance *c*^*q*^ should give rise to the minimal error. This shift selection process allows for the identification of penetrance from pseudo-bulk data without having access to single-cell data for novel perturbations.

Note that such shift selections are unavoidable to obtain accurate predictions, as the unseen perturbation could have very distinct penetrance. However, the performance of the model will still rely on whether SCCVAE successfully learns the underlying regulatory information, as attributing a single scalar parameter alone will not lift sufficient capacity to generalize to unseen perturbations.

#### 2.2.1 Computational complexity

Consider *N*_*I*_ perturbations at inference time. By generating *B* cells per perturbation, the computational complexity of evaluating *C* possible shift values is *O*(*N*_*I*_ *BCm*), where *m* is the number of genes in the considered expression data. The typical choice of *B* is around 32 *∼* 128. A finer level of shift selection with larger *C* results in more accurate results, where the computational complexity scales linearly in *C*.

## 3 Results

We evaluate on the single-gene perturbational datasets by [31] which contains normalized gene expression transcripts of essential genes for two different cell lines, K562 and RPE1 cells, where *m*=8563 on K562 cells and *m*=8749 on RPE1 cells.

In our first set of analyses, we focus on the cancerous K562 cell line and examine a subset of perturbations that induced distinct expression distributions from the control distribution, in order to reduce the noise to signal ratio. This subset is chosen by first filtering out the perturbations with less than 200 single cells. Out of the remaining perturbations, we consider only the perturbations that yield an average five-fold cross validation score of greater than 0.6 in a logistic regression task distinguishing a perturbed cell from negative controls. This ensures that the subset of perturbations (*N* = 279) are distinct from control and require nontrivial predictions. Further details can be found in Appendix A.3.1.

We additionally include experiments on the entire K562 cell line (without filtering perturbations) by comparison, along with experiments on a selected subset of the RPE1 cell line. The additional experimental results can be found in Appendix A.3.6

### 3.1 Setup

#### 3.1.1 In-distribution experiments

In this set of experiments, we evaluate how well different models can interpolate observed perturbations. All cells in the test set are drawn from the same perturbational distributions as those in the training set. For each perturbation in the dataset, we assign 70% of the cells to the training set, 10% to the validation set, and the remaining 20% to the test set.

#### 3.1.2. Out-of-distribution experiments

In this set of experiments, we evaluate the performance of different models on generalizing to unseen perturbations. We designate the train/test split so that there is no overlap between the test set and the train set perturbations. First, we randomly select 20% of the perturbations to make up the test set. Out of the remaining perturbations, we assign 85% of the cells from each distribution to the training set, and the remaining 15% to the validation set. To account for variation in the amount of distributional shift that occurs from random selection of test perturbations, we repeat this experiment using five different train/test splits, so that each perturbation is included in the test set for one of the splits. Results are averaged across all splits. For each unseen perturbation *X*^*q*^ (defined in Section 2.2), we select *c*^*q*^ by searching through the range of all learned *c*^*p*^ in the model and select the value that minimizes mean squared error for pseudo-bulk predictions. Details of this search process are found in Appendix A.1.3.

In both in-distribution and out-of-distribution tasks, training hyperparameters were selected based on minimizing loss (defined in Section 2.1) within the validation set, drawn from the same distribution as the training set and determined as described above. Hyperparameters are selected through the validation set separately for each train/test split. As such, no points from the test set are seen in the hyperparameter search for any given split. Further details are given in Appendix A.1.

### 3.2 Prediction of Single-Cell Expressions

To evaluate SCCVAE, we consider six metrics: Mean squared error, Pearson correlation, maximum mean discrepancy, energy distance, and fraction of genes changed/unchanged from control in the opposite direction of ground truth. Formulas for computing each metric are found in Appendix A.2. Each perturbation is evaluated separately, and we report the mean and standard deviation across all test perturbations for each metric. Each metric is computed on the set of all essential genes and on the more difficult task of predicting just the top 50 most highly variable genes.

#### 3.2.1 SCCVAE versus single-cell-level baselines

Here we focus on perturbational effects at the single-cell level. For baselines, we compare against the control cell distribution, a popular deep learning model (GEARS [12]), and a transformer-based model (scGPT [19]) for single-cell predictions.

Results of the in-distribution experiments can be found in Table 3 in Appendix A.3.2, where we observed that SCCVAE outperforms or is on par with both baselines on all metrics. This shows the benefit of utilizing an expressive network in SCCVAE to interpolate observed perturbations.

Table 1 shows the results of the out-of-distribution experiments. It can be seen that the SCCVAE outperforms both the GEARS and control baselines for generalizing to unseen perturbations. The improvements above baselines for single-cell based metrics, i.e., MMD and Energy Distance, are more pronounced when evaluating on all essential genes. This is further illustrated in Fig. 2, where we observe that SCCVAE matches the perturbation distributions much better than the baselines. The fraction of perturbations changed or in the same direction from control is also improved in the SCCVAE predictions. SCCVAE additionally achieves better MSE and Pearson R than the baselines, although all baselines perform well over these metrics. However, in SCCVAE there is much less variation across different perturbations.

**Table 1:** SCCVAE vs baselines on the out-of-distribution task. Results are averaged across five different splits, each containing a different set of perturbations in the test set covering the entire set of all perturbations. Both when evaluating all essential genes and top 50 genes, SCCVAE achieves better metrics than control or GEARS for every metric except Pearson correlation, where all three methods achieve similar mean values but SCCVAE has lower variance in its predictions. SCCVAE outperforms a transformer-based model, scGPT, on both distributional-based metrics. When compared to a simple linear model from Ahlmann-Eltze et al [20], SCCVAE achieves similar metrics to the linear model, while additionally achieving low distributional loss, expanding its scope beyond the linear model.

| All genes | Control | GEARS | scGPT | Linear | SCCVAE |
| --- | --- | --- | --- | --- | --- |
| MSE | 0.00817±0.00575 | 0.00734±0.00524 | 0.0101±0.00105 | **0.00438±0.000392** | 0.00542±0.002 |
| PearsonR | 0.989±0.00748 | 0.989±0.00738 | **0.997±0.000644** | 0.992±0.00071 | 0.992±0.004 |
| MMD | 0.240±0.0391 | 0.239±0.0405 | 0.266±0.0104 | - | **0.229±0.0193** |
| Energy Dist. | 0.0363±0.0138 | 0.0333±0.0132 | 0.0337±0.00929 | - | **0.0299±0.00958** |
| Frac. Same | 0.496±0.0175 | 0.551±0.0590 | **0.717±0.0468** | 0.627±0.0123 | 0.630±0.0509 |
| Frac. Changed | 0.471±0.0175 | 0.449±0.0590 | **0.283±0.0468** | 0.373±0.0123 | 0.370±0.0509 |

Table 1: SCCVAE vs baselines on the out-of-distribution task.
| Top 50 genes | Control | GEARS | scGPT | Linear | SCCVAE |
| --- | --- | --- | --- | --- | --- |
| MSE | 0.00969±0.00830 | 0.00879±0.00770 | 0.0215±0.00362 | **0.00561±0.000359** | 0.00651±0.00537 |
| PearsonR | 0.995±0.00503 | 0.995±0.00498 | **0.998±0.000742** | 0.996±0.000303 | 0.996±0.00376 |
| MMD | 0.246±0.0512 | 0.245±0.0531 | 0.313±0.0246 | - | **0.236±0.0323** |
| Energy Dist. | 0.0385±0.0199 | 0.0372±0.0203 | 0.0588±0.0221 | - | **0.0338±0.0160** |
| Frac. Same | 0.492±0.0541 | 0.553±0.0945 | **0.721±0.0763** | 0.618±0.0163 | 0.612±0.0820 |
| Frac. Changed | 0.468±0.0541 | 0.447±0.0945 | **0.279±0.0763** | 0.382±0.0163 | 0.388±0.0821 |

**Figure 2.**
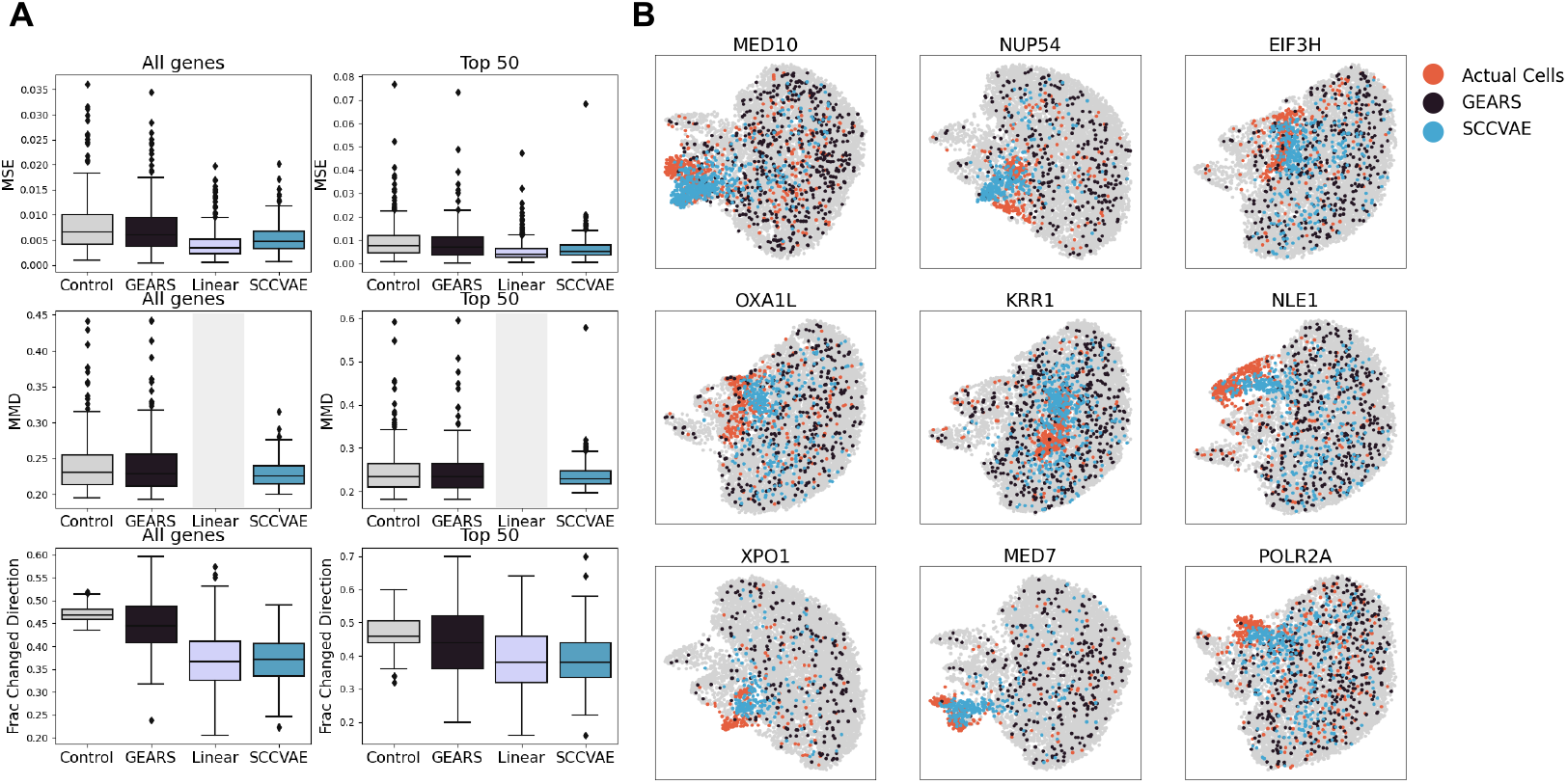
(A) When comparing quantitative performance on the OOD task of GEARS vs SCCVAE to the control distribution, GEARS on average learns the control distribution but SCCVAE is more closely able to approximate the ground truth and is comparable to the linear method across all metrics. This effect is very pronounced when observing all essential genes, in the case of top 50 genes there are a few outlier perturbations with unusually high error. (B) SCCVAE and GEARS UMAP visualizations versus ground-truth perturbations on select perturbations in the OOD task. Consistent with quantitative results, GEARS outputs match the control distribution while SCCVAE outputs match the perturbationally distinct ground truth.

Additionally, although scGPT attains better Pearson correlations and fraction changed/same metrics, the distributional error metrics (MMD and Energy Distance) are all higher than control. This is indicative of the ability of transformer-based models to capture large-scale patterns in the data better than the causal model, but the large distributional error indicates that the model still overfits and cannot generalize to finer distributional details, unlike in the causal model.

#### 3.2.2. SCCVAE versus linear model

The performance of the SCCVAE model in comparison to the simple linear model explored in Ahlmann-Eltze et. al is showcased in Table 1 above. ([20]). We use the principal component based representation for the perturbations, same as SCCVAE. It can be seen that both the SCCVAE and the linear model achieve on par performance for average bulk metrics. Recall our discussion in Section 2.2; notably, the linear model is able to predict the bulk expression well without explicitly specifying the penetrance. This may be because unseen perturbations in this dataset have similar penetrance to those in the training set–specifically, their guides have similar efficiencies. As such, one possible alternative for SCCVAE to perform shift selection in this library can be using predicted bulk expression from the linear model.

On the other hand, the SCCVAE is able to handle single-cell data on a distributional level, while the linear model is limited to bulk analysis. Extending the linear model to capture a distribution over gene expressions is challenging, as it requires specifying higher-order moments of the distribution (beyond just the mean) and modifying the loss function to account for these moments.

### 3.3 Ablation Studies

We evaluate the SCCVAE architecture using different choices of graph *G* (Section 2.1). The results in Table 1 are generated using a learned DAG without sparsity restrictions. We compare the performance using a non-hybrid model with a conditional variational autoencoder (Conditional), a model using a pre-specified gene regulatory network inferred using a causal discovery algorithm on negative control cells [32] (Causal-GSP), and a model using a pre-specified randomly generated graph (Random). We evaluate the performance in the out-of-distribution setting. The details of this setup are described in Appendix A.1.

The results of this comparison are shown in Table 2 and Figure 3. We observe that SCCVAE with a learned graph or an inferred DAG identified by GSP outperform the conditional model and the random graph models. Because bulk-level metrics experience minimal change, we focus on MMD. These results further demonstrate that incorporating a mechanistic causal model improves the model performance on generalizing to unseen perturbations.

**Table 2:** Ablation studies. SCCVAE with a learned graph (SCCVAE) consistently achieves superior MSE and MMD values Both the conditional model and the pre-specified graph-based model (Causal-GSP) still outperform the GEARS baselines from Table 1, and the pre-specified graph-based model outperforms the conditional model, but both are more restrictive than SCCVAE with a learned graph. The models with random causal graphs achieve high error, as is expected, but outperform all other models on the fraction same/changed metrics. This is likely due to the shifts being tuned to extreme values during the training/shift selection process to account for the poor distributional match from the random graph.

| All genes | SCCVAE | Conditional | Causal-GSP | Random |
| --- | --- | --- | --- | --- |
| MSE | **0.00542**±0.00295 | 0.00580±0.00399 | 0.00569±0.00367 | 0.00779±0.00270 |
| PearsonR | **0.992**±0.00469 | 0.991±0.00552 | 0.991±0.00532 | 0.991±0.00432 |
| MMD | **0.229**±0.0193 | 0.232±0.0293 | 0.231±0.0271 | 0.268±0.0259 |
| Energy Dist. | **0.0299**±0.00958 | **0.0299**±0.0112 | 0.0300±0.0104 | 0.0332±0.00703 |
| Frac. Same | 0.630±0.0509 | 0.618±0.0655 | 0.622±0.0630 | **0.644**±0.0521 |
| Frac. Changed | 0.370±0.0509 | 0.382±0.0655 | 0.378±0.0630 | **0.356**±0.0521 |

Table 2: Ablation studies. SCCVAE with a learned graph (SCCVAE) consistently achieves superior MSE and MMD values.
| Top 50 genes | SCCVAE | Conditional | Causal-GSP | Random |
| --- | --- | --- | --- | --- |
| MSE | **0.00651**±0.00537 | 0.00728±0.00685 | 0.00689±0.00641 | 0.0101±0.00542 |
| PearsonR | **0.996**±0.00376 | **0.996**±0.00436 | **0.996**±0.00425 | 0.995±0.00377 |
| MMD | **0.236**±0.0323 | 0.243±0.0435 | 0.240±0.0413 | 0.289±0.0413 |
| Energy Dist. | 0.0338±0.0160 | 0.0329±0.0182 | **0.0328**±0.0167 | 0.0389±0.013 |
| Frac. Same | 0.612±0.082 | 0.610±0.0939 | 0.616±0.0912 | **0.633**±0.0717 |
| Frac. Changed | 0.388±0.0821 | 0.390±0.0938 | 0.384±0.0912 | **0.367**±0.0716 |

**Table 3:** In-distribution results for SCCVAE compared to GEARS and the control baseline, for all essential genes and for top 50 highly variable genes, averaged across all perturbational distributions.

| All genes | Control | GEARS | SCCVAE |
| --- | --- | --- | --- |
| MSE | 0.0131±0.00641 | 0.0118±0.00568 | **0.0114±0.004** |
| PearsonR | **0.982±0.00841** | **0.982±0.00833** | **0.982±0.007** |
| MMD | 0.242±0.0431 | 0.239±0.0440 | **0.220±0.01** |
| Energy Distance | 0.0363±0.0200 | 0.0325±0.0176 | **0.0296±0.01** |
| Fraction Same | 0.497±0.0127 | 0.541±0.0319 | **0.587±0.04** |
| Fraction Changed | 0.471±0.0127 | 0.459±0.0319 | **0.413±0.04** |

| Top 50 genes | Control | GEARS | SCCVAE |
| --- | --- | --- | --- |
| MSE | 0.0153±0.00970 | 0.0134±0.00846 | **0.0126±0.006** |
| PearsonR | **0.991±0.00595** | **0.991±0.00565** | **0.991±0.004** |
| MMD | 0.249±0.0602 | 0.244±0.0583 | **0.227±0.03** |
| Energy Distance | 0.0425±0.0336 | 0.0370±0.0317 | **0.0316±0.02** |
| Fraction Same | 0.501±0.0589 | 0.548±0.0762 | **0.586±0.08** |
| Fraction Changed | 0.459±0.0589 | 0.452±0.0762 | **0.414±0.08** |

**Figure 3.**
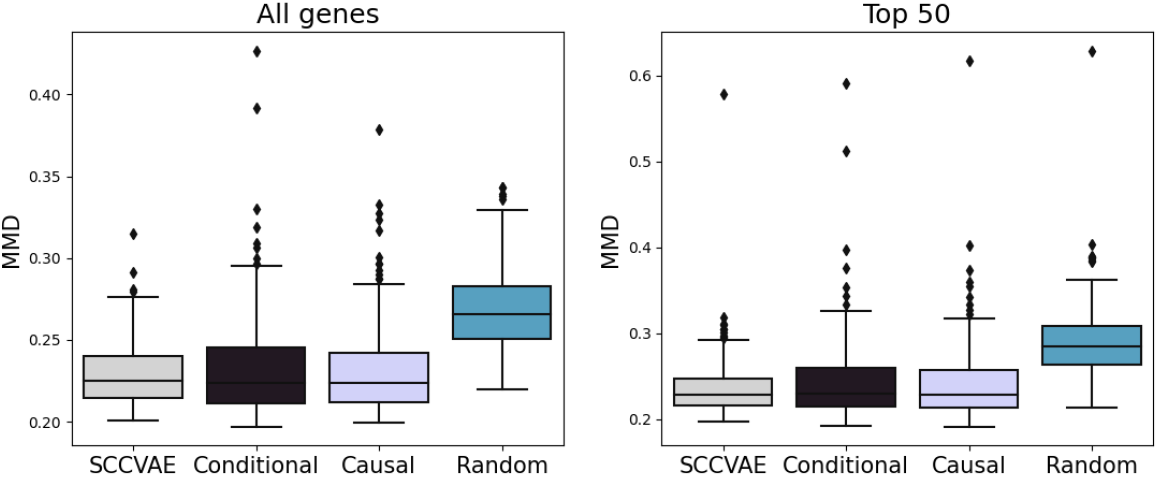
Distribituional loss (MMD) on SCCVAE ablations. The learned causal graph in the SCCVAE model achieves lower MMD than conditional, sparse causal graph, and random graph equivalents.

#### 3.3.1 SCCVAE versus conditional model

The causal SCCVAE achieves lower error than the equivalent conditional model, where the input to the decoder is 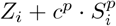. The difference is most noticeable in the distributional loss metrics (MMD and Energy Distance), as the conditional model has no way of learning interactions between latent variables that aid in out-of-distribution generalization.

#### 3.3.2 Learned versus pre-specified causal graphs

Our results indicate that the sparse graph learned by GSP is overly restrictive in the OOD setting. Overall, it does not generalize as well to the out-of-distribution split as SCCVAE with a learned graph, in terms of the distributional-level metrics (MMD and Energy Distance). However, it is notable that Causal-GSP still performs consistently better than the conditional model, especially on top 50 variable genes, indicating that the sparse graph identified by the GSP algorithm still successfully learns certain regulatory relationships.

#### 3.3.3. SCCVAE versus random DAGs

The error metrics (MSE, Pearson, MMD, Energy Distance) for the random graph model are significantly worse than the other models. However, the average direction of change metrics are improved, likely because the shift value *c* (defined in Section 2.2) is selected to overcompensate for the lack of distributional precision in the predictions. Recall that the shift selection process for *c* is a required step for out-of-distribution prediction in our mechanistic model, because the penetrance for novel genetic perturbations is unknown. However, when the mechanisms in the model are mis-specified (e.g., with a random graph), shift selection is unable to correct the prediction. This provides further evidence that the ability of model to generalize to unseen perturbations depends on whether the regulatory information can be successfully learned.

### 3.4 Latent Gene Embeddings

To examine the latent representations learned by SCCVAE, we first investigate if they contain information on whether the perturbations induce distribution shifts from the negative controls. Figure 4 compares the MMD between *X*^*p*^ and *X*^Ø^ in the gene expression space with the *L*_2_ distance between *U* ^*p*^ and *U* ^Ø^ for each *p* in the latent space in all five OOD splits. For each split, these two distances are highly correlated, which is consistent with the knowledge that SCCVAE learns the perturbation changes through *U* ^*p*^, since *U* ^*p*^ = (I_*n*_ *− A*)^*−*1^(*Z* + *c*^*p*^ *· S*^*p*^) and *U* ^Ø^ = (I_*n*_ *− A*)^*−*1^*Z* (Section 2.1) A further breakdown of the scatter plots into training and testing perturbations is found in Appendix A.3.3.

**Figure 4.**
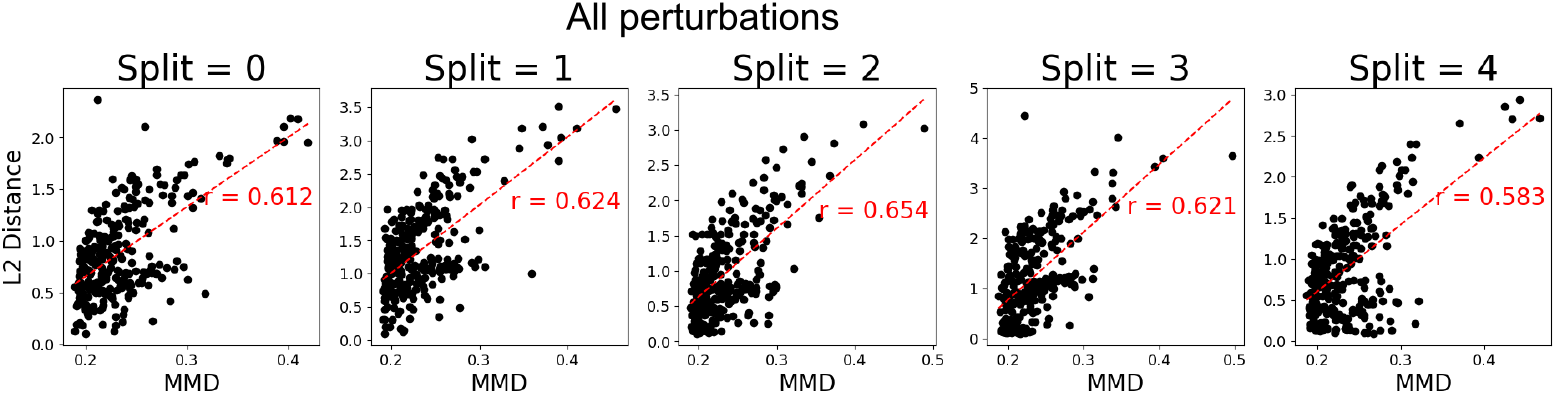
Distance from the control distribution in the observational space vs the latent space (*U* ^*p*^), for all perturbations in each OOD split. The Euclidean distance in the latent space is strongly correlated with the MMD in the expression space.

By modulating *c*^*p*^, we can simulate varying penetrance of the perturbation, pushing the post perturbational distribution closer or further away from control. Figure 5 demonstrates this in action on the gene MED7, as *c*^*p*^ is set to various values in [-1, 3]. As *c* is increased from -1 to 2, the error decreases until reaching a minimum at *c ≈* 2.0 (Fig. 5A). Likewise, the output distribution matches the control distribution when *c*^*p*^ *≈* 0, most closely matches the ground truth in the dataset at *c*^*p*^ *≈* 2.0, but can continue to be extended beyond that value as values of *c*^*p*^ *>* 2.0 push the prediction further away from the control distribution (Fig. 5B). Further analysis of the effect of shift selection on other perturbation predictions is given in Appendix A.3.4.

**Figure 5.**
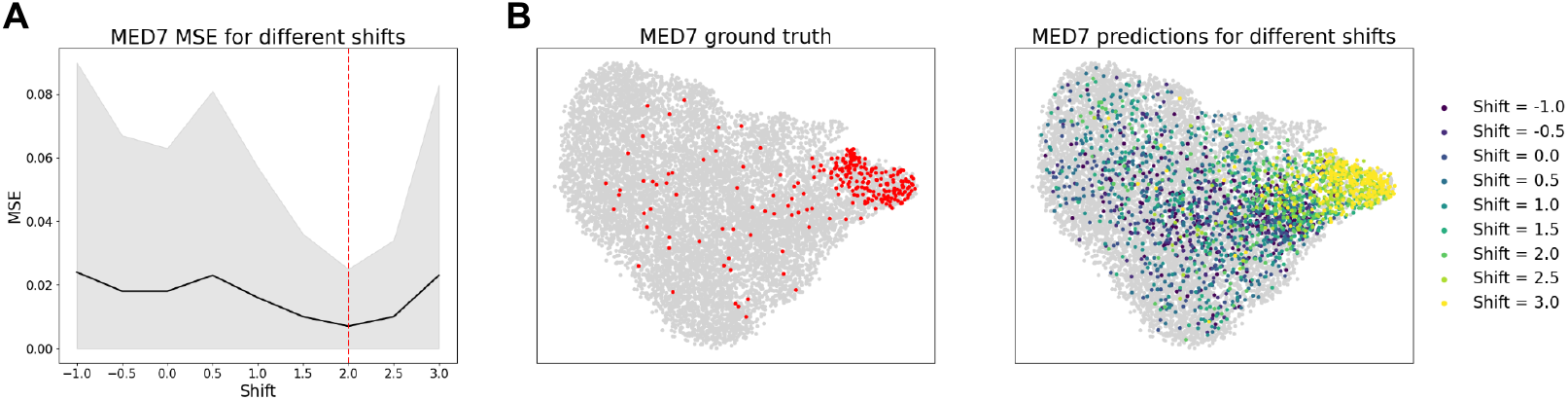
(A) For OOD test perturbations, the shift is selected to minimize MSE of pseudo-bulk predictions. (B) Shift values closer to zero result in output predictions closer to control, and larger magnitude shift values result in more distinct perturbational distributions. The shift value that most closely matches the ground truth is *c ≈* 2.0.

Figure 6 visualizes *U*^*p*^ embeddings for both training and testing distributions from one of the out-of-distribution splits. These embeddings were generated by encoding the perturbations (*p* or *q*) along with a sample of control cells (*X*), and no reparameterization was used to simulate the effect of taking the average after infinite samples. It can be seen that perturbations of genes with similar functions map to similar variables in the latent space, and identifiable perturbational modules (such as mediator complex genes, ribosomal proteins, cell cycle regulators, proteasome subunits, and genes involved in DNA replication and repair) are clustered together in the latent space, regardless of if the genes in the module were included in the original training set or not. As such, it is possible to infer the function of an unknown gene using SCCVAE by examining the functions of its neighbors when encoded to *U* ^*p*^. More visualizations of perturbation modules in *U* ^*p*^ can be found in Appendix A.3.5.

**Figure 6.**
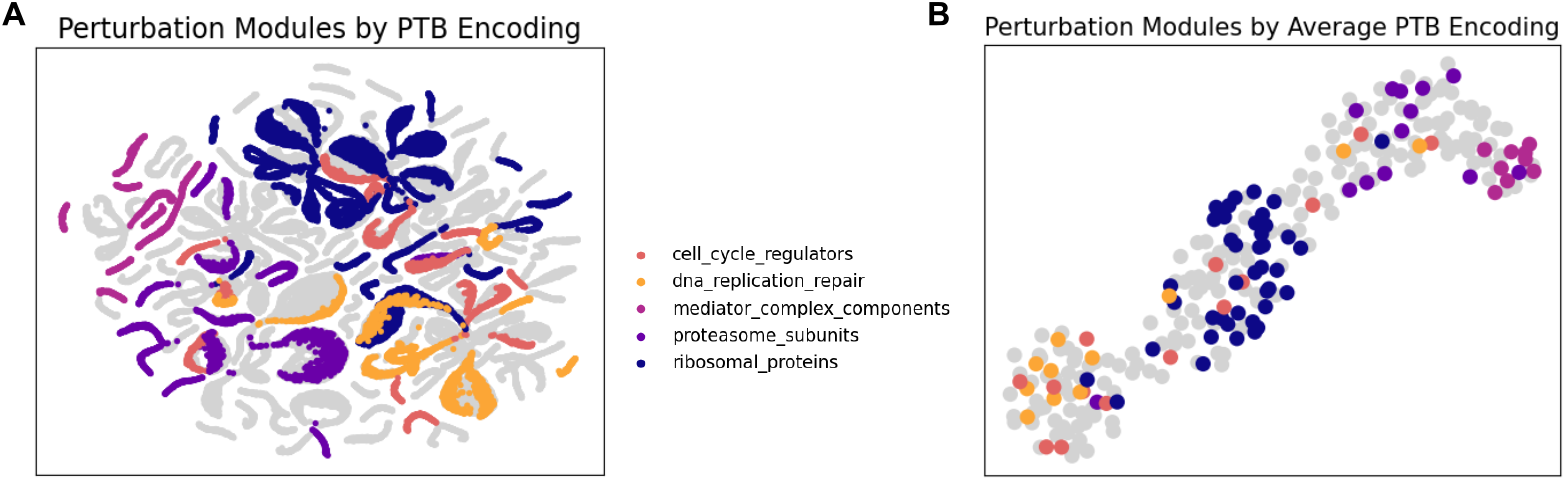
(A) UMAP Visualizations of latent causal variables with various functional perturbation modules. Genes belonging to the same perturbation module are close together within the latent space. (B) When visualizing just the average *U* ^*p*^ of each perturbation, the perturbation modules form distinct clusters in the latent space.

## 4 Conclusions

In this work, we demonstrated the ability of a hybrid variational causal model (SCCVAE) to effectively model gene expression outcomes following single-gene perturbations, particularly in scenarios when extrapolating to unseen perturbations. By leveraging a learned mechanistic model within a flexible variational neural network, SCCVAE is able to both interpolate observed perturbations and generalize to unseen perturbations. These findings underscore the importance of modeling gene regulatory relationships in the predictive model for accurate predictions of genetic interventions. Additionally, the ability of SCCVAE to learn shifts through a structural causal model provides a robust foundation for simulating gene knockdown experiments with varying penetrance.

Our findings open various directions for future exploration. First, we used SCCVAE for single-gene knockdown perturbations. Extending our model to different types of perturbations, or different combinations of perturbations, is an interesting future direction. Second, the current version of SCCVAE implicitly uses a Gaussian likelihood in the output space (i.e., *P* (*X*|*Z*) is modeled as a Gaussian distribution) to approximate the normalized gene expressions. However, raw gene expressions can be modeled using a zero-inflated negative binomial (ZINB) distribution, the parameters of which reveal additional gene-specific information about the perturbed distribution. Extending and further analyzing SCCVAE on ZINB based likelihood is of interest. Lastly, we demonstrated how to incorporate a mechanistic model into a variational autoencoder framework. It would be interesting to build upon this work to test incorporation with alternative generative models.

## Acknowledgements

We would like to acknowledge E. Forte for manuscript proofreading. E.L. was supported by the Eric and Wendy Schmidt Center at the Broad Institute. J.Z. was partially supported by an Apple AI/ML PhD Fellowship. C.U. was partially supported by NCCIH/NIH (1DP2AT012345), ONR (N00014-22-1-2116 and N00014-24-1-2687), the United States Department of Energy (DE-SC0023187), the Eric and Wendy Schmidt Center at the Broad Institute, and a Simons Investigator Award.

## A Supplementary Information

### A.1 Code and Data Availability

Code and relevant data for the SCCVAE model and experiments are available at https://github.com/uhlerlab/sccvae.

#### A.1.1. Architecture and training

The SCCVAE model is composed of the following:

1. *Expression encoder* : After the input cell expressions are log normalized, they are passed through 2 fully connected hidden layers of size 1024. Mean and variance encoders are fully connected layers of output size 512 that accept the output of the shared encoding. Each intermediate layer uses a leaky ReLU activation function with negative slope 0.2.

2. *Shift encoder* : 1 fully connected hidden layer of size 1024, followed by Leaky ReLU activation (negative slope=0.2) and an output layer of size 512. Output is softmaxed.

3. *SCM* : The causal graph is simulated using a size 512 *×* 512 parameter initialized to a Gaussian distribution with mean 0 and variance 0.1, multiplied by an upper triangular binary mask. Shift values are initialized as a 1 *×* 512 model parameter.

4. *Expression decoder* : 2 fully connected hidden layers of size 1024, followed by an output layer of size 8563. All layers except for output use LeakyReLU with negative slope 0.2.

The model was trained using a learning rate of 5e-4, a linear KL annealing schedule with *β*_*max*_ = 1, and MMD scale *γ* = 10. To represent the perturbation *p*, the top 50 PCA components were used. To stabilize training, a reconstruction term for the control distribution is added, where *U* ^Ø^ is decoded with *c*^*p*^ = 0 to reconstruct *X*^Ø^. This term is scaled by 0.25 for in-distribution splits and out-of-distribution splits 0, 2, 3, and 4, and scaled by 1 for out-of-distribution split 1. Inclusion of this term reduces the risk of exploding gradients, but does not significantly affect numerical results.

#### A.1.2 Hyperparameter selection

Hyperparameter selection for learning rate, *β, γ*, and neural network size were performed using a grid search and selected based on lowest validation error according to the validation sets described in Section 3.1. Hyperparameters were selected from the following lists of possible values:

|  |  |
| --- | --- |
| Learning Rate | [1e-3, 5e-4, 1e-4] |
| $\beta$ | [1, 2, 4, 8] |
| $\gamma$ | [1, 2, 4, 8, 10, 15] |
| X reconstruction term | [0.0, 0.25, 0.5, 0.75, 1.0] |
| Number of layers in encoder/decoder | [1, 2, 3] |
| Layer size | [512, 1024] |

#### A.1.3 Shift selection

After training, when presented with an unseen perturbation, we search for an optimal shift value within the range of *c*^*p*^ learned by the model. Using increments of 0.1, we grid search every possible shift value within range. For each shift value, we substitute for *c*^*q*^ and compute 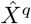. The final value of *c*^*q*^ is set to the value that minimizes 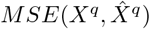, comparing to the mean expression value obtained from bulk data for perturbation *q*. Since we do not have access to an alterative bulk screen, we utilize pseudo-bulk expressions computed from the same dataset.

### A.2 Quantitative Metrics

#### A.2.1 Mean Squared Error

The mean squared error (MSE) metric evaluates the error between the means of the predicted and ground-truth distributions:

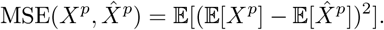

The final value reported is averaged across all genes of the mean normalized expression level.

#### A.2.2 Pearson Correlation

We also evaluate the Pearson correlation between the distribution means, using a similar approach to the evaluation of MSE:

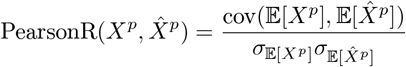

where cov(*·*) is the covariance and *σ*_(*·*)_ is the standard deviation of each mean expression value.

#### A.2.3. Maximum Mean Discrepancy

The maximum mean discrepancy (MMD) evaluates a distribution level error:

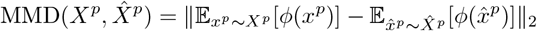

where *ϕ*(*·*) is the Gaussian kernel function.

#### A.2.4. Energy Distance

To compute energy distance (*D*^2^), we compare the prediction and ground truth on a distributional level as follows:

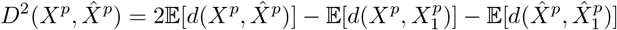

where 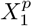 and 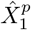 are independent samples from the same distributions as *X*^*p*^ and 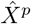 respectively (implemented as random permutations of cells). The distance function is given by *d*(*X, Y*) = 𝔼 [(*X−Y*)^2^].

To correct for the effect of distribution sample size on error magnitude, MMD and Energy distance were computed in batches of size 32 averaged across the entire test dataset.

#### A.2.5 Direction of Changes

We evaluate the quality of the model predictions on gene differential expression. For each perturbation, we compare the fraction of genes where the direction of change from the control cell distribution is the same as that of the ground truth (Fraction same), as well as the fraction of genes that are changed in the opposite direction (Fraction changed):

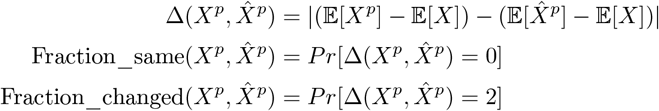

where *X* is the control control cell distribution.

### A.3 Additional Results

#### A.3.1 Subset of perturbations

Figure 7 shows the process for selecting perturbations for training. From the starting set of perturbations, approximately 20% are perturbations with 200 cells or more. Out of these, around 68% have a logistic regression score of 0.6 or more. Then, we remove perturbations that have no identified causal relationships with other observed genes, or are not included in the expression vector. This leaves 279 perturbational distributions for our analysis.

**Figure 7.**
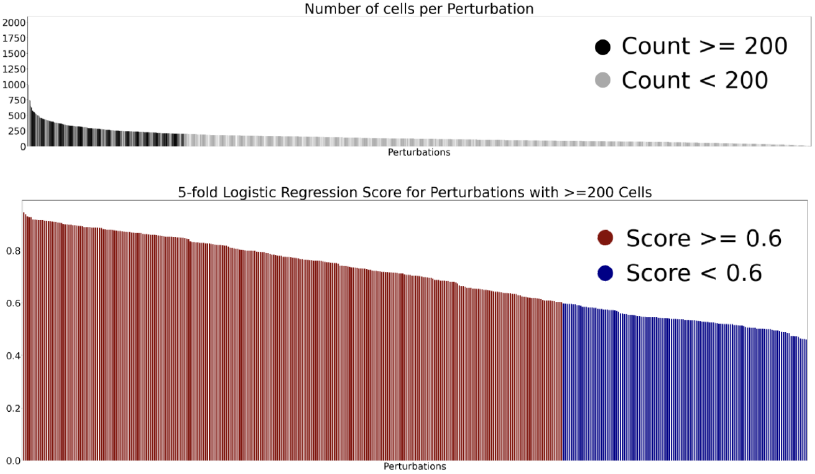
(A) We limit our analysis to perturbations with over 200 single cells. (B) Logistic regression filtering for perturbational distributions on the remaining perturbations. Only perturbations wtih a 5-fold cross validation logistic regression score of 0.6 or higher are considered.

#### A.3.2 In-distribution results

Table 3 shows the results of SCCVAE training on the in-distribution data set. It can be seen that SCCVAE beats both GEARS and control baselines for all metrics, both on the entire set of essential genes and on the top 50 most highly variable genes in the control data set. In Table 4, we observe similar trends to the OOD results in that the original SCCVAE model outperforms the other versions of the model in MSE and MMD metrics. However, we note here that the Causal-GSP model does not outperform the conditional model (except for energy distance), unlike in the OOD task, and the conditional model is also able to outperform the original SCCVAE on the energy distance metric. This is because the ability to utilize a causal graph and generalize to new cells is less crucial for the in-distribution task, since the test cells are from the same distribution as train cells.

**Table 4:**
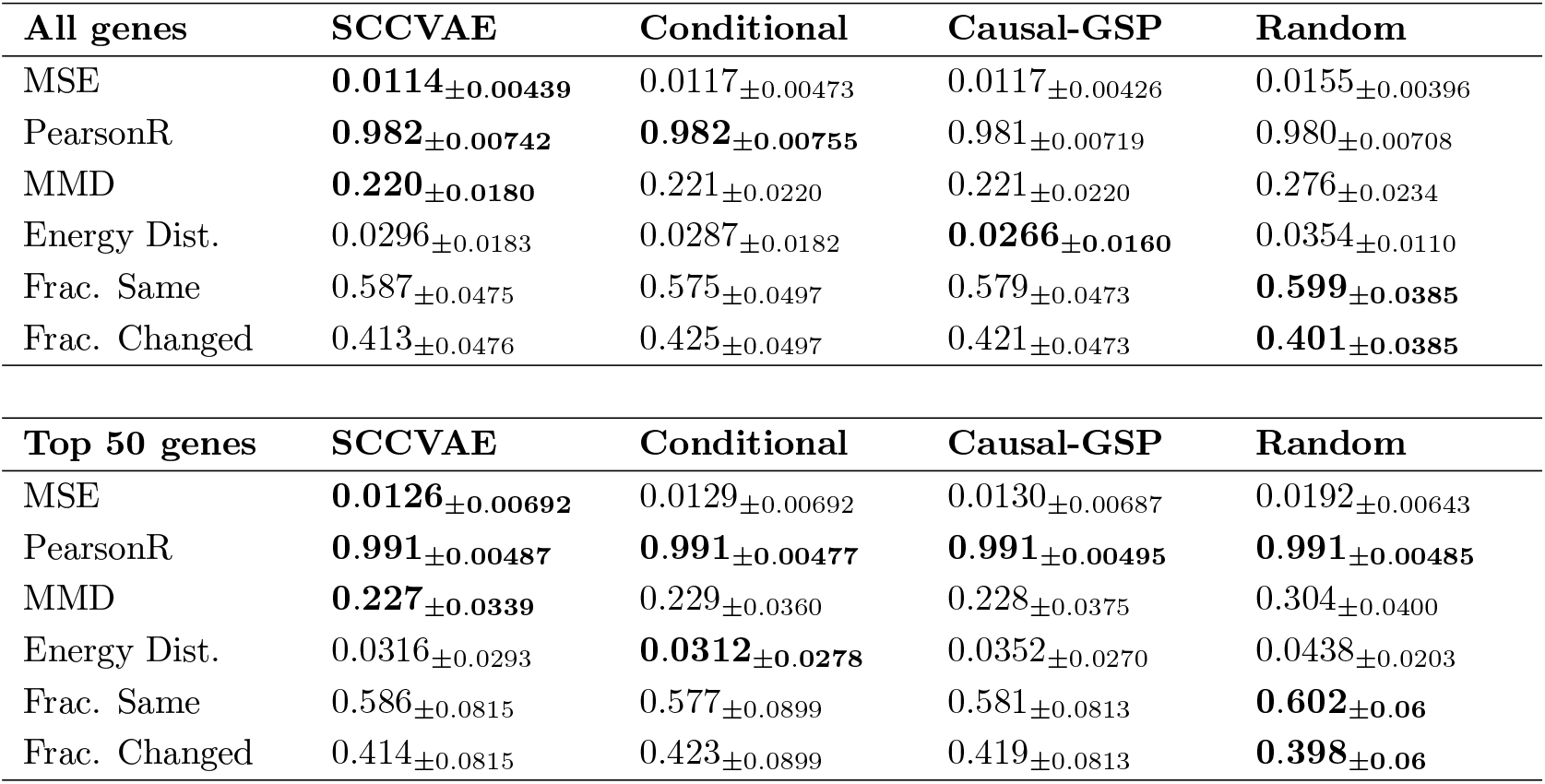
Ablation studies on the in-distribution task. Unlike in the OOD task, the conditional and causal GSP models achieve better performance than the original SCCVAE for the energy distance metric.

#### A.3.3 Breakdown of distance scatterplots by training/testing distributions

Figure 8 looks at the correlation between MMD and latent space distance (similar to Figure 4) for train and test perturbations separately. There is a strong positive correlation in training test perturbations, as expected, but the test set perturbations also exhibit positive correlations across all five splits.

**Figure 8.**
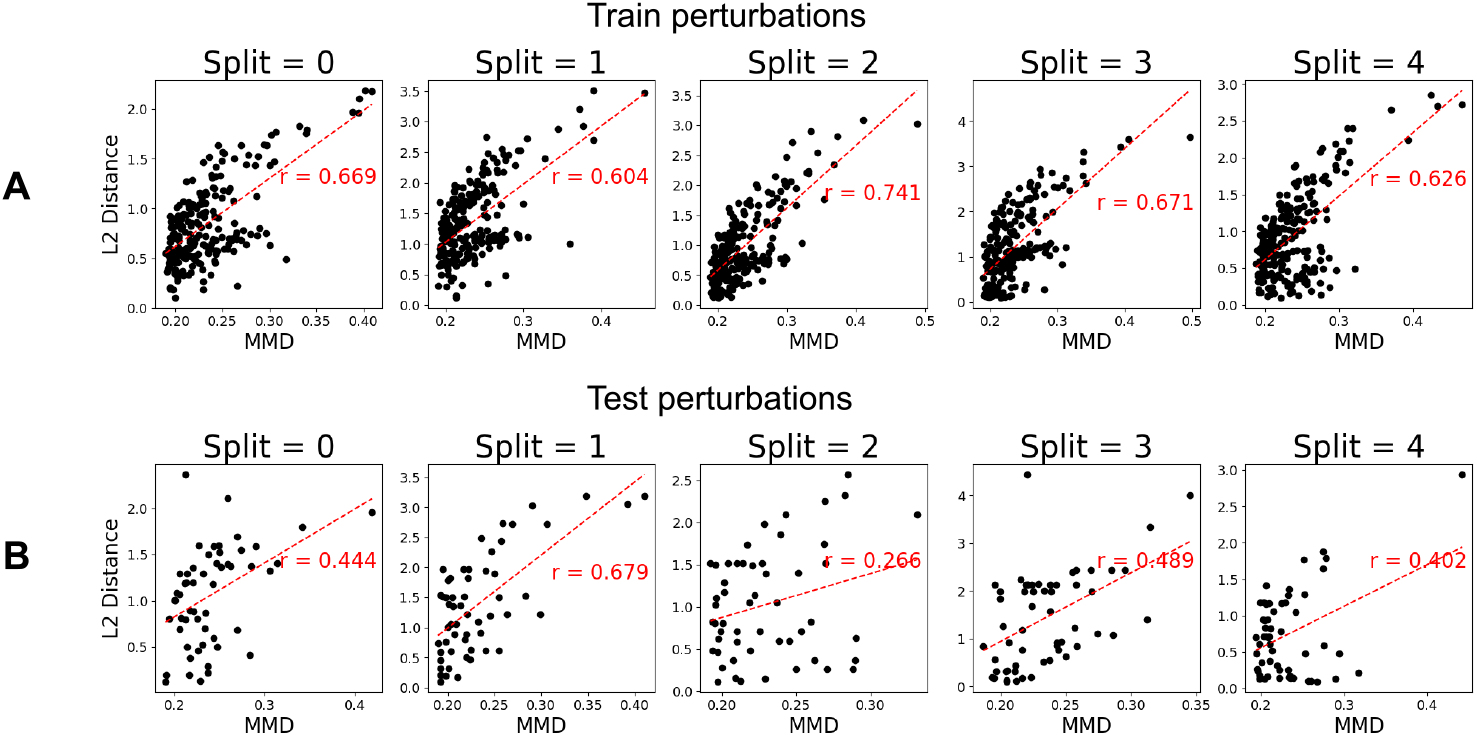
Continuation of Figure 4. (A) In just the training/validation set perturbations, the latent variables are learned directly during training (since no shift selection is required). This leads to stronger correlations overall. (B) In the test set, latent variables are computed after selecting shift to minimize MSE. Although the correlation is weaker than the training set perturbations, it is consistently positive across all splits.

#### A.3.4 Additional shift selection results

Figure 9 shows the same procedure as Figure 5 repeated on a different gene, NLE1. For NLE1, the post-perturbational expression distribution is much closer to the control distribution, and the shift value that yield the lowest error is also much smaller in magnitude (*c*^*p*^ *≈* 0.5). We can see that as *c*^*p*^ become larger, the predicted distribution also moves farther away from the ground truth.

**Figure 9.**
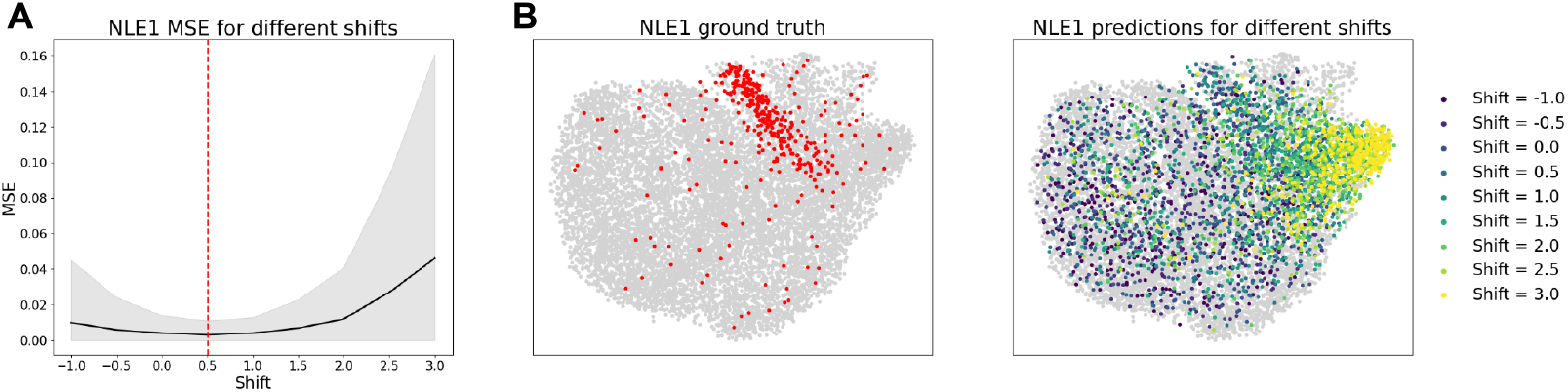
(A) Shift selection for a different perturbation (NLE1), where the optimal shift value is closer to control. (B) When *c*^*p*^ is increased, we can visualize the postperturbational cell expression distribution under a higher intensity gene knockdown.

#### A.3.5 Additional perturbation modules

Figure 10 visualizes the embeddings of 12 different functional perturbation modules in *U* ^*p*^ latent space. As stated already, the embeddings of genes within one perturbational module form a cluster in the latent space. We observe that even across different perturbation modules affecting the same cellular functions, similar functional modules are also grouped closely in the latent space. For example, all genes encoding ribosomal proteins or ribosome biogenesis are close to each other, chromatin and DNA controlling genes are grouped closely together, and splicing genes are also grouped near each other.

**Figure 10.**
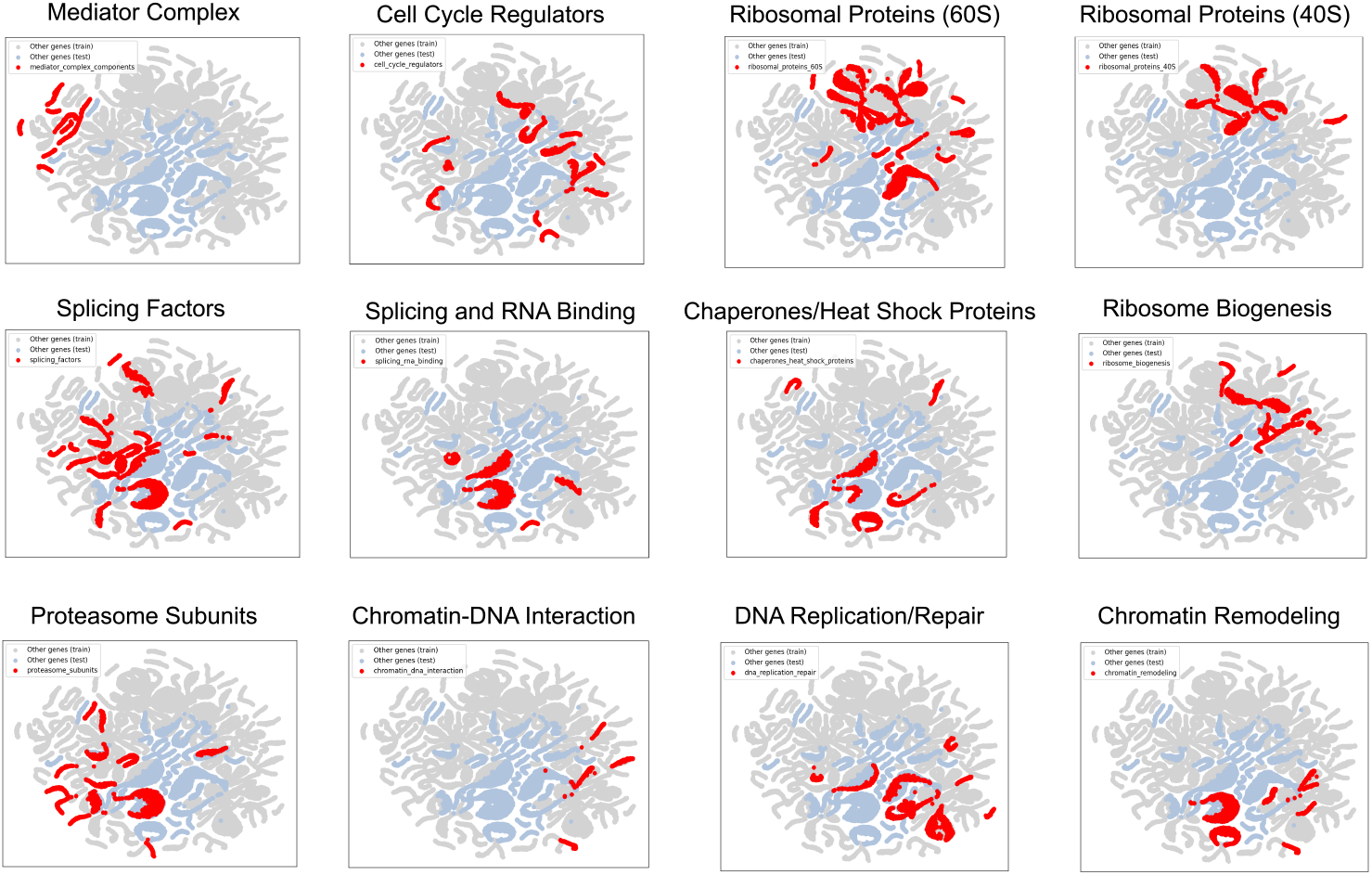
Individual perturbation modules. In each figure, the red regions indicate *U* ^*p*^ embeddings of cells that are part of the perturbation module, the grey regions indicate *U* ^*p*^ embeddings of non-module cells from the training set, and the blue regions indicate *U* ^*q*^ embeddings of non-module cells from the test set.

#### A.3.6 Interpretation of the learned SCM

Figure 11 visualizes the structural causal model learned by SCCVAE (Section 2.1) in the in-distribution experiment on *N* = 279 perturbations on K562 cells. As the learned graph *𝒢* is encoded by the adjacency matrix *A* ∈ ℝ ^512*×*512^, we first use *A* to read out a learned network. The perturbations alter variables in this network by the encoded shift vectors 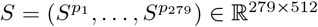.

**Figure 11.**
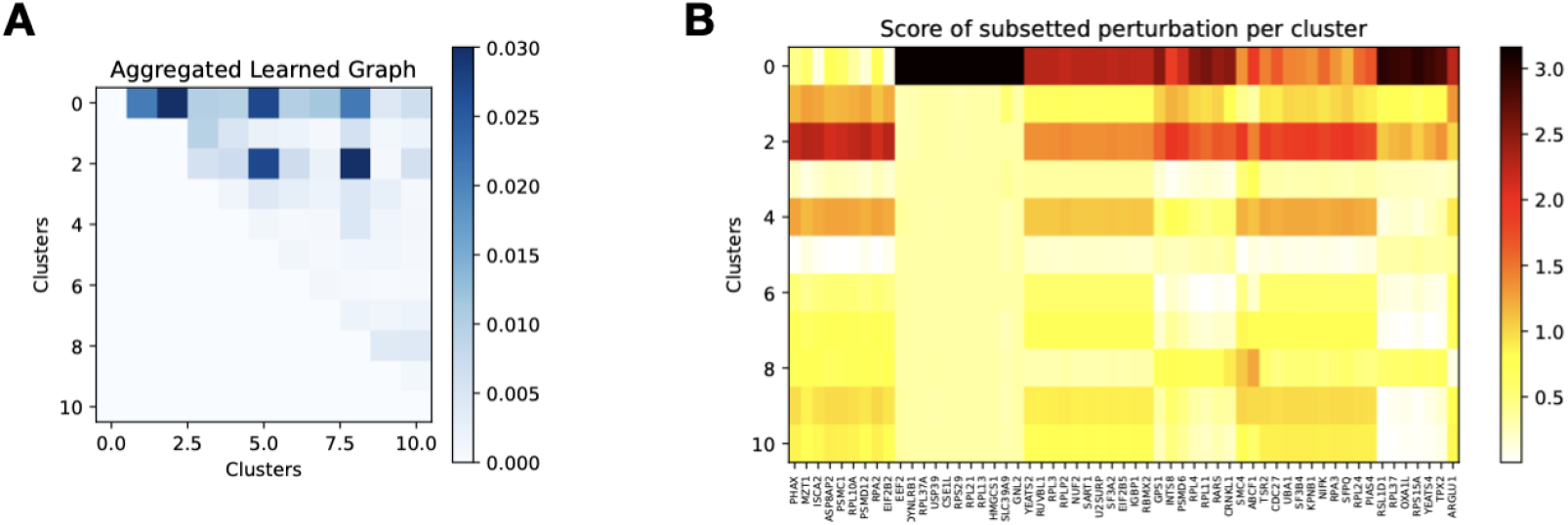
SCM Learned by SCCVAE. (A) Learned adjacency matrix *A* aggregated by clusters found using the encoded shift vectors *S*. (B) Learned shift vectors *S* averaged using the clusters found in (A) on a subset of perturbations.

To interpret the learned network, we cluster the dimensions of this network according to their responses to the perturbations. In particular, we use K-means clustering based on *S*^*⊤*^ with 11 clusters. We then aggregate the adjacency matrix *A* by averaging the scores between dimensions in each pair of clusters. A cluster is considered to be more upstream than a distinct cluster if in less than half of the entries in the sub-matrix of *A* corresponding these two clusters are zero. By re-orienting the clusters from upstream to downstream, we visualize the aggregated absolute connections in this network in Figure 11 (A).

To interpret how each perturbation *S*^*p*^ alters the individual clusters in the aggregated network, we average the entries of *S*^*p*^ corresponding to each cluster and use the averaged vector as the scoring for this perturbation. We cluster the perturbations based on the averaged scoring vector and visualize 10 random perturbations per cluster in Figure 11 (B). Notably cluster 0 is mainly contributed by the Ribosome family. A deeper understanding of these clusters may reveal biological insights.

#### A.3.7 Comparison across datasets

We demonstrate the capacity of SCCVAE over other datasets. Out of the considered baselines, GEARS has shown to have consistent performances across metrics, therefore we compare against GEARS for analysis over other datasets. In Table 5, we compare SCCVAE to GEARS and control cell baselines on the entire dataset, without selecting for high signal-to-noise ratio perturbational distributions. Here, we observe that SCCVAE still generally out-performs baselines, although the difference is less dramatic than in the selected perturbations case. Additionally, both GEARS and control baselines appear to achieve lower error on the full dataset compared to the selected dataset, whereas SCCVAE exhibits the opposite trend and achieves higher error on the full dataset. The most likely explanation for this is that the full dataset contains many perturbational distributions that are very similar to the control distribution. As a result, the control baselines are lower, and GEARS, which outputs similar predictions to control, achieves lower error. SCCVAE, which tends to identify differences from the control distribution, generally performs better when the perturbed distribution is distinct from the control distribution.

**Table 5:** Evaluation on the entire K562 single-cell dataset, without selecting for high signal-to-noise ratio perturbations. Since GEARS exhibited the best performance out of the two model baselines, we evaluate against GEARS and the control baseline. SCCVAE achieves lower error than control or GEARS for all metrics except MMD with all essential genes and all metrics except MMD and Energy Distance with top 50 genes. Compared to evaluation on the selected perturbation dataset, evaluation on all genes appears to be an easier task for all models. This is likely due to the similarity of most distributions to control, as evidenced by the fact that GEARS exhibits a larger drop in performance on the selected dataset than SCCVAE.

| All genes | Control | GEARS | SCCVAE |
| --- | --- | --- | --- |
| MSE | 0.0077±0.0077 | 0.00731±0.00745 | **0.00657±0.004** |
| PearsonR | 0.989±0.0109 | 0.989±0.0107 | **0.99±0.006** |
| MMD | 0.213±0.0286 | **0.212±0.0308** | 0.217±0.02 |
| Energy Distance | 0.0272±0.0138 | 0.0264±0.0136 | **0.0262±0.01** |
| Fraction Same | 0.484±0.0067 | 0.522±0.0226 | **0.544±0.04** |
| Fraction Changed | 0.483±0.0067 | 0.478±0.0226 | **0.456±0.04** |

Table 5: Evaluation on the entire K562 single-cell dataset, without selecting for high signal-to-noise ratio perturbations.
| Top 50 genes | Control | GEARS | SCCVAE |
| --- | --- | --- | --- |
| MSE | 0.00861±0.00962 | 0.00809±0.0093 | **0.00739±0.006** |
| PearsonR | **0.995±0.00643** | **0.995±0.00634** | **0.995±0.004** |
| MMD | 0.215±0.0446 | **0.214±0.044** | 0.221±0.03 |
| Energy Distance | 0.0291±0.0243 | **0.0279±0.0237** | 0.0283±0.0 |
| Fraction Same | 0.52±0.0722 | 0.528±0.0744 | **0.548±0.08** |
| Fraction Changed | 0.48±0.0722 | 0.472±0.0744 | **0.452±0.08** |

Table 6 shows results upon evaluating a selected RPE1 dataset, also from Replogle et al. [31]. While SCCVAE continues to outperform GEARS and control cell baselines on most metrics when evaluating on all essential genes, both GEARS and SCCVAE consistently do not beat the control cell baseline when only the top 50 highly variable genes are considered, with SCCVAE usually achieving lower error than GEARS. There are two possible causes for this phenomenon. First, the RPE1 dataset is smaller than the K562 dataset, and the neural network based methods may require more data than is available to learn optimally. Second, the RPE1 cell line is non-cancerous, while the K562 cell line is cancerous, meaning that they derive from fundamentally different distributions. It is possible that these models are more performant on cancerous cell lines, as they may better capture the underlying biological variability and gene expression patterns that are more pronounced in cancerous cells due to their altered regulatory mechanisms and higher levels of genetic instability. This suggests that these models might implicitly exploit features or patterns specific to cancer biology, leading to diminished performance on non-cancerous cell lines like RPE1, which exhibit more stable and less variable gene expression profiles. Future work could explore tailoring these models to account for differences in cell line characteristics or incorporating stronger priors to improve generalization across multiple different cell lines.

**Table 6:**
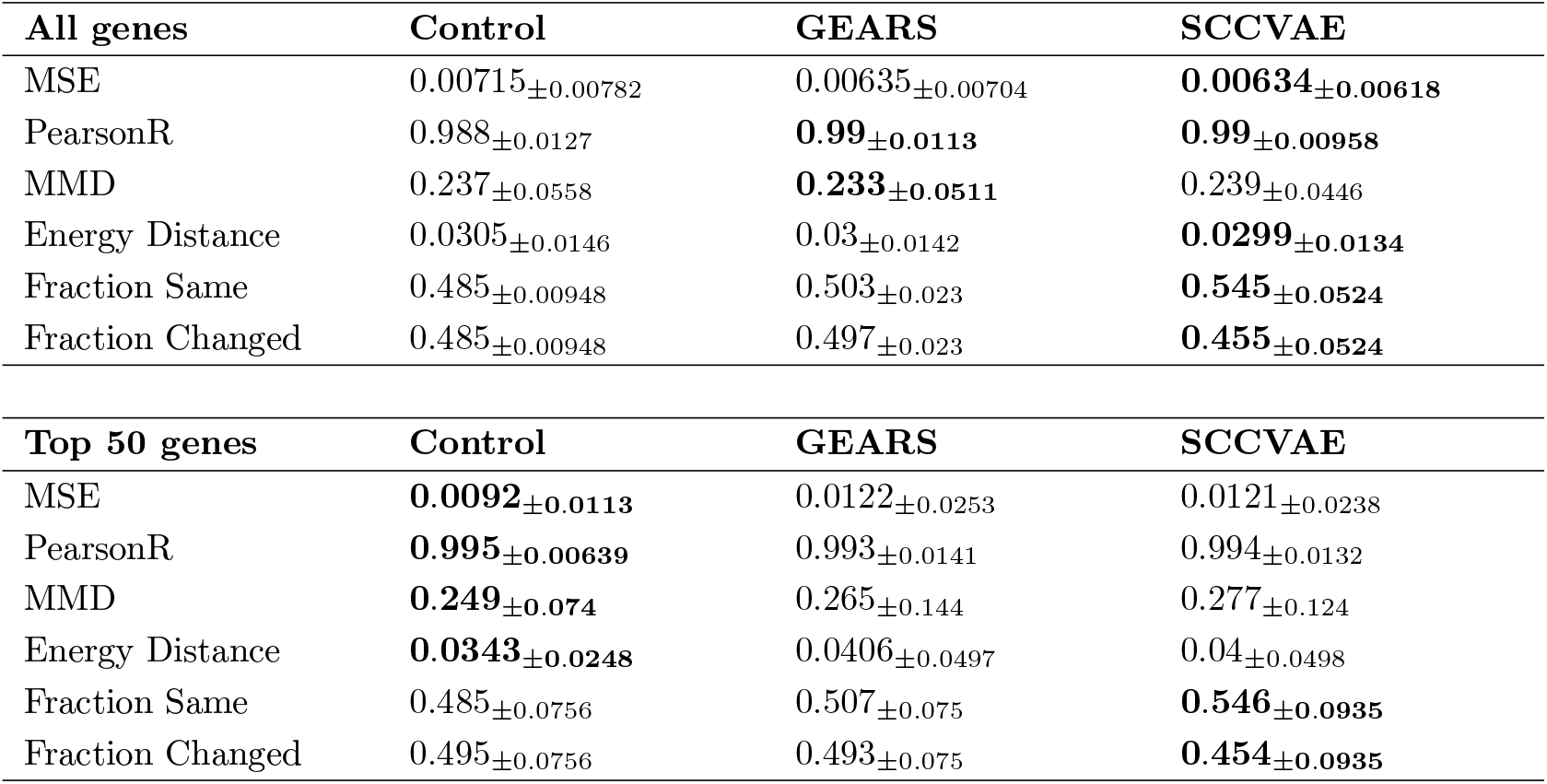
Evaluation on the RPE1 cell line. SCCVAE achieves lower error than baselines when considering all essential genes, but the control cell baseline has lower error than both GEARS and SCCVAE when considering the top 50 highly variable genes.

## Notes

### Competing Interest Statement

The authors have declared no competing interest.

## References

[1] Christoph Bock, Paul Datlinger, Florence Chardon, Matthew A Coelho, Matthew B Dong, Keith A Lawson, Tian Lu, Laetitia Maroc, Thomas M Norman, Bicna Song, et al. High-content crispr screening. Nature Reviews Methods Primers, 2(1):1–23, 2022.

[2] Jeremy D Grevet, Xianjiang Lan, Nicole Hamagami, Christopher R Edwards, Laavanya Sankaranarayanan, Xinjun Ji, Saurabh K Bhardwaj, Carolyne J Face, David F Posocco, Osheiza Abdulmalik, et al. Domain-focused crispr screen identifies hri as a fetal hemoglobin regulator in human erythroid cells. Science, 361(6399):285–290, 2018.

[3] Weiwei Cheng, Shaopeng Wang, Zhe Zhang, David W Morgens, Lindsey R Hayes, Soojin Lee, Bede Portz, Yongzhi Xie, Baotram V Nguyen, Michael S Haney, et al. Crispr-cas9 screens identify the rna helicase ddx3x as a repressor of c9orf72 (ggggcc) n repeat-associated non-aug translation. Neuron, 104(5):885–898, 2019.

[4] Gary Dixon, Heng Pan, Dapeng Yang, Bess P Rosen, Therande Jashari, Nipun Verma, Julian Pulecio, Inbal Caspi, Kihyun Lee, Stephanie Stransky, et al. Qser1 protects dna methylation valleys from de novo methylation. Science, 372(6538):eabd0875, 2021.

[5] Tim Wang, Jenny J Wei, David M Sabatini, and Eric S Lander. Genetic screens in human cells using the crispr-cas9 system. Science, 343(6166):80–84, 2014.

[6] Traver Hart, Megha Chandrashekhar, Michael Aregger, Zachary Steinhart, Kevin R Brown, Graham MacLeod, Monika Mis, Michal Zimmermann, Amelie Fradet-Turcotte, Song Sun, et al. High-resolution crispr screens reveal fitness genes and genotype-specific cancer liabilities. Cell, 163(6):1515–1526, 2015.

[7] Atray Dixit, Oren Parnas, Biyu Li, Jenny Chen, Charles P Fulco, Livnat Jerby-Arnon, Nemanja D Marjanovic, Danielle Dionne, Tyler Burks, Raktima Raychowdhury, et al. Perturb-seq: dissecting molecular circuits with scalable single-cell rna profiling of pooled genetic screens. cell, 167(7):1853–1866, 2016.

[8] Britt Adamson, Thomas M Norman, Marco Jost, Min Y Cho, James K Nuñez, Yuwen Chen, Jacqueline E Villalta, Luke A Gilbert, Max A Horlbeck, Marco Y Hein, et al. A multiplexed single-cell crispr screening platform enables systematic dissection of the unfolded protein response. Cell, 167(7):1867–1882, 2016.

[9] Joseph M Replogle, Reuben A Saunders, Angela N Pogson, Jeffrey A Hussmann, Alexander Lenail, Alina Guna, Lauren Mascibroda, Eric J Wagner, Karen Adelman, Gila Lithwick-Yanai, et al. Mapping information-rich genotype-phenotype landscapes with genome-scale perturb-seq. Cell, 185(14):2559–2575, 2022.

[10] Mohammad Lotfollahi, Anna Klimovskaia Susmelj, Carlo De Donno, Leon Hetzel, Yuge Ji, Ignacio L Ibarra, Sanjay R Srivatsan, Mohsen Naghipourfar, Riza M Daza, Beth Martin, et al. Predicting cellular responses to complex perturbations in high-throughput screens. Molecular systems biology, 19(6):e11517, 2023.

[11] Hengshi Yu and Joshua D Welch. Perturbnet predicts single-cell responses to unseen chemical and genetic perturbations. BioRxiv, pages 2022–07, 2022.

[12] Y Roohani, K Huang, and J. Leskovec. Predicting transcriptional outcomes of novel multigene perturbations with GEARS. In Nat Biotechnol 42, 2024.

[13] Romain Lopez, Natasa Tagasovska, Stephen Ra, Kyunghyun Cho, Jonathan Pritchard, and Aviv Regev. Learning causal representations of single cells via sparse mechanism shift modeling. In Conference on Causal Learning and Reasoning, pages 662–691. PMLR, 2023.

[14] Yulun Wu, Robert A Barton, Zichen Wang, Vassilis N Ioannidis, Carlo De Donno, Layne C Price, Luis F Voloch, and George Karypis. Predicting cellular responses with variational causal inference and refined relational information. arXiv preprint arXiv:2210.00116, 2022.

[15] Hananeh Aliee, Ferdinand Kapl, Soroor Hediyeh-Zadeh, and Fabian J Theis. Conditionally invariant representation learning for disentangling cellular heterogeneity. arXiv preprint arXiv:2307.00558, 2023.

[16] Xinming Tu, Jan-Christian Hutter, Zitong Jerry Wang, Takamasa Kudo, Aviv Regev, and Romain Lopez. A supervised contrastive framework for learning disentangled representations of cell perturbation data. bioRxiv, pages 2024–01, 2024.

[17] Jennifer E Rood, Anna Hupalowska, and Aviv Regev. Toward a foundation model of causal cell and tissue biology with a perturbation cell and tissue atlas. Cell, 187(17):4520–4545, 2024.

[18] Christina V Theodoris, Ling Xiao, Anant Chopra, Mark D Chaffin, Zeina R Al Sayed, Matthew C Hill, Helene Mantineo, Elizabeth M Brydon, Zexian Zeng, X Shirley Liu, et al. Transfer learning enables predictions in network biology. Nature, 618(7965):616–624, 2023.

[19] Haotian Cui, Chloe Wang, Hassaan Maan, Kuan Pang, Fengning Luo, Nan Duan, and Bo Wang. scgpt: toward building a foundation model for single-cell multi-omics using generative ai. Nature Methods, pages 1–11, 2024.

[20] Constantin Ahlmann-Eltze, Wolfgang Huber, and Simon Anders. Deep learning-based predictions of gene perturbation effects do not yet outperform simple linear methods. BioRxiv, pages 2024–09, 2024.

[21] Charlotte Bunne, Andreas Krause, and Marco Cuturi. Supervised training of conditional monge maps. Advances in Neural Information Processing Systems, 35:6859–6872, 2022.

[22] Jiaqi Zhang, Kristjan Greenewald, Chandler Squires, Akash Srivastava, Karthikeyan Shanmugam, and Caroline Uhler. Identifiability guarantees for causal disentanglement from soft interventions. Advances in Neural Information Processing Systems, 36, 2024.

[23] Thomas Gaudelet, Alice Del Vecchio, Eli M Carrami, Juliana Cudini, Chantriolnt-Andreas Kapourani, Caroline Uhler, and Lindsay Edwards. Season combinatorial intervention predictions with salt & peper. arXiv preprint arXiv:2404.16907, 2024.

[24] Kenji Kamimoto, Blerta Stringa, Christy M Hoffmann, Kunal Jindal, Lilianna Solnica-Krezel, and Samantha A Morris. Dissecting cell identity via network inference and in silico gene perturbation. Nature, 614(7949):742–751, 2023.

[25] Jiaqi Zhang, Louis Cammarata, Chandler Squires, Themistoklis P Sapsis, and Caroline Uhler. Active learning for optimal intervention design in causal models. Nature Machine Intelligence, 5 (10):1066–1075, 2023.

[26] Payam Dibaeinia and Saurabh Sinha. Sergio: a single-cell expression simulator guided by gene regulatory networks. Cell systems, 11(3):252–271, 2020.

[27] Peter Spirtes, Clark Glymour, and Richard Scheines. Causation, prediction, and search. MIT press, 2001.

[28] Judea Pearl. Causality. Cambridge university press, 2009.

[29] Diederik P Kingma. Auto-encoding variational bayes. arXiv preprint arXiv:1312.6114, 2013.

[30] Arthur Gretton, Karsten M Borgwardt, Malte J Rasch, Bernhard Schölkopf, and Alexander Smola. A kernel two-sample test. The Journal of Machine Learning Research, 13(1):723–773, 2012.

[31] Joseph M. Replogle, Reuben A. Saunders, Angela N. Pogson, Jeffrey A. Hussmann, Alexander Lenail, Alina Guna, Lauren Mascibroda, Eric J. Wagner, Karen Adelman, Jessica L. Bonnar, Marco Jost, Thomas M. Norman, and Jonathan S. Weissman. Mapping information-rich genotype-phenotype landscapes with genome-scale perturb-seq. Cell, 2022.

[32] Liam Solus, Yuhao Wang, and Caroline Uhler. Consistency guarantees for greedy permutation-based causal inference algorithms. Biometrika, 108(4):795–814, 2021.

